# RNA structure inference through chemical mapping after accidental or intentional mutations

**DOI:** 10.1101/169953

**Authors:** Clarence Y. Cheng, Wipapat Kladwang, Joseph Yesselman, Rhiju Das

**Affiliations:** Department of Biochemistry, Stanford University, Stanford, CA 94305, United States; Department of Physics, Stanford University, Stanford, CA 94305, United States

## Abstract

Despite the critical roles RNA structures play in regulating gene expression, sequencing-based methods for experimentally determining RNA base pairs have remained inaccurate. Here, we describe a multidimensional chemical mapping method called M2-seq (mutate-and-map read out through next-generation sequencing) that takes advantage of sparsely mutated nucleotides to induce structural perturbations at partner nucleotides and then detects these events through dimethyl sulfate (DMS) probing and mutational profiling. In special cases, fortuitous errors introduced during DNA template preparation and RNA transcription are sufficient to give M2-seq helix signatures; these signals were previously overlooked or mistaken for correlated double DMS events. When mutations are enhanced through error-prone PCR, *in vitro* M2-seq experimentally resolves 33 of 68 helices in diverse structured RNAs including ribozyme domains, riboswitch aptamers, and viral RNA domains with a single false positive. These inferences do not require energy minimization algorithms and can be made by either direct visual inspection or by a new neural-net-inspired algorithm called M2-net. Measurements on the P4-P6 domain of the *Tetrahymena* group I ribozyme embedded in *Xenopus* egg extract demonstrate the ability of M2-seq to detect RNA helices in a complex biological environment.

**SIGNIFICANCE STATEMENT:** The intricate structures of RNA molecules are crucial to their biological functions but have been difficult to accurately characterize. Multidimensional chemical mapping methods improve accuracy but have so far involved painstaking experiments and reliance on secondary structure prediction software. A methodology called M2-seq now lifts these limitations. Mechanistic studies clarify the origin of serendipitous M2-seq-like signals that were recently discovered but not correctly explained and also provide mutational strategies that enable robust M2-seq for new RNA transcripts. The method detects dozens of Watson-Crick helices across diverse RNA folds *in vitro* and within frog egg extract, with low false positive rate (< 5%). M2-seq opens a route to unbiased discovery of RNA structures *in vitro* and beyond.

## INTRODUCTION

Inference of RNA structures using experimental data is a crucial step in understanding RNA’s biological functions throughout living organisms. Chemical mapping methods have the potential to reveal RNA structural features *in situ* by probing which nucleotides are protected from attack by chemical modifiers. The resulting experimental data can be used as pseudo-energies to guide secondary structure modeling by computational algorithms, raising the prospect of transcriptome-wide RNA structure determination (1, 2).

Despite these advances, the accuracy of RNA structure inference approach through chemical mapping and sequencing remains under question (3-8). For example, models of the 9 kb HIV-1 RNA genome have been repeatedly revised with updates to the selective 2′-OH acylation by primer extension (SHAPE) protocol, data processing, and computational assumptions (2, 9-11), and the majority of this RNA’s helices remain uncertain. Even for small RNA domains, SHAPE and dimethyl sulfate (DMS; methylation of N1 and N3 atoms at A and C) have produced misleading secondary structures for ribosomal domains and blind modeling challenges that have been falsified through crystallography or mutagenesis (3, 7) (12, 13). In alternative approaches based on photo-activated crosslinkers, many and perhaps the majority of helix detections appear to be false positives, based on ribosome data *in vitro* and *in vivo* (14, 15).

The confidence and structural accuracy of chemical mapping methods can be improved by applying perturbations to the RNA sequence prior to chemical modification. In the mutate-and-map strategy, mapping not just the target RNA sequence but also a comprehensive library of point mutants reveals which nucleotides respond to perturbations at every other nucleotide, enabling direct inference of pairs of residues that interact to form structure (16, 17). The resulting models have been consistently accurate at nucleotide resolution in RNA-puzzles and other blind tests for riboswitches and ribozymes solved by crystallography, with helix recovery rates of >90% and false positive rates under 10%, with errors typically involving minor register shifts or edge base pairs (2, 18). However, the mutate-and-map approach has required synthesis and parallel mapping of many mutant RNAs and, so far, has only been applied to RNAs under 200 nucleotides in length probed *in vitro.*

Here we introduce <u>m</u>utate-and-<u>m</u>ap read out by next-generation <u>seq</u>uencing (M2-seq), which carries out RNA preparation, mutation, and mapping in a one-pot experiment. Tests on ribozyme domains, viral domains, and riboswitch aptamers that form diverse RNA structures evaluate the ability of M2-seq to detect Watson-Crick base pairs *in vitro*, with signals that can be confirmed through visual inspection. We introduce a simple algorithm M2-net that automatically recovers these helices with a low false positive rate (< 5%) and without register shifts that have been previously problematic for chemical mapping approaches. As a proof-of-concept for more complex biological experiments, we demonstrate direct detection of the majority of helices in the P4-P6 domain of the group I *Tetrahymena* ribozyme embedded in biologically active eukaryotic cell extract, and describe prospects for further applications in RNA structural biology.

## RESULTS

### Workflow of mutate-and-map read out through next-generation sequencing

The M2-seq workflow tested herein is summarized in Figure 1. First, DNA templates were prepared from PCR assembly (short constructs) or PCR from plasmids (long constructs). To ensure mutate-and-map signals, we prepared samples with a low frequency (~10^−3^ per nucleotide) of additional mutations as described previously (16) or using error-prone PCR (19). We also prepared samples without additional mutations to probe unexplained data correlations observed in recent high-DMS experiments (2, 20). Then, we transcribed RNAs from these DNA pools, prepared them into the desired state (e.g. Mg^2+^-induced folding *in vitro*) and modified the RNA with DMS. Reverse transcription was performed under mutational profiling conditions (with Superscript II and Mn^2+^) to install mutations into cDNAs across from DMS modifications (21). The full-length cDNAs were amplified by PCR, and the resulting libraries were sequenced by paired-end Illumina sequencing. An initial M2-seq map was generated by recording the positions of all the correlated mutations. The data were displayed in a two-dimensional heat-map visualization analogous to that used for prior mutate-and-map experiments: a 1D chemical mapping profile was estimated for each single-nucleotide-variant in the RNA, each profile was normalized by the total number of reads with a mutation at that position, and the profiles were stacked according to the mutation. As described below and in SI Results, a more sophisticated analysis is possible that attempts to separate mutations based on their expected source (e.g., those installed during library preparation vs. those introduced later by reverse transcription across from chemical modifications). However, we will mainly describe results with a simple mutation-counting approach, which provides an initial unbiased visualization.

**Figure 1.**
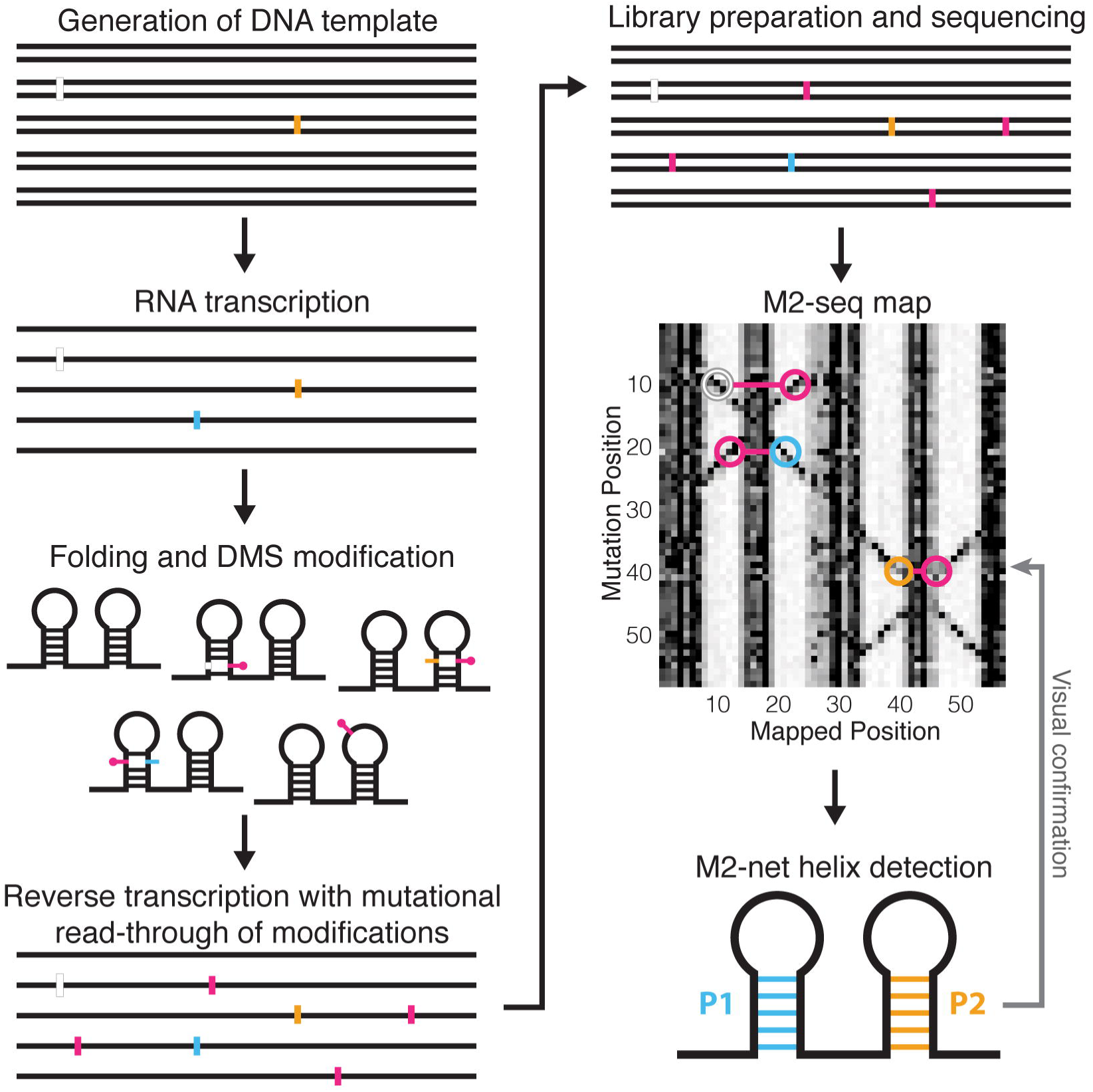
Workflow for M2-seq, mutate-and-map read out by next-generation sequencing. DNA template is generated by PCR assembly of oligonucleotides, PCR from a plasmid, or error-prone PCR potentially introducing deletions (white) or mutations (gold). Then, RNA is transcribed (potentially introducing further mutations, cyan), folded, and DMS-modified. Reverse transcription is performed under conditions that favor mutational read-through of DMS-modified nucleotides, recording those positions as mutations (magenta), and cDNAs are PCR-amplified to generate double-stranded DNA library. Libraries are subjected to next-generation sequencing, and resulting reads are analyzed by demultiplexing, alignment to reference sequences, and correlated mutation counting to generate a mutate-and-map-seq (M2-seq) dataset (simulated here). Doublestranded RNA helices give rise to cross-diagonal features in these maps that can be automatically recognized by M2-net and confirmed visually.

### Mutational profiling provides precise M2-seq information in a single-pot experiment

We first confirmed that applying the mutational profiling readout to single-mutant libraries would give secondary structure signals similar to capillary electrophoresis-based M2, which relies on reverse transcription termination at modified residues rather than mutational read-through. For this comparison, we investigated the P4-P6 domain of the 158-nucleotide *Tetrahymena* group I ribozyme (Fig. 2A), a widely used model system for tests of RNA chemical mapping methods (2, 3, 22). In addition, we prepared DNA templates for the wild-type and 158 point mutants of each nucleotide to its complement (16) and then pooled these molecules prior to the transcription step, so that all subsequent steps could be carried out in one tube. The M2-seq data for this initial pooled-mutation experiment are shown in Fig. 2B, after applying the pipeline described in Fig. 1 to generate 1D chemical mapping reactivity profiles for each mutation position.

**Figure 2.**
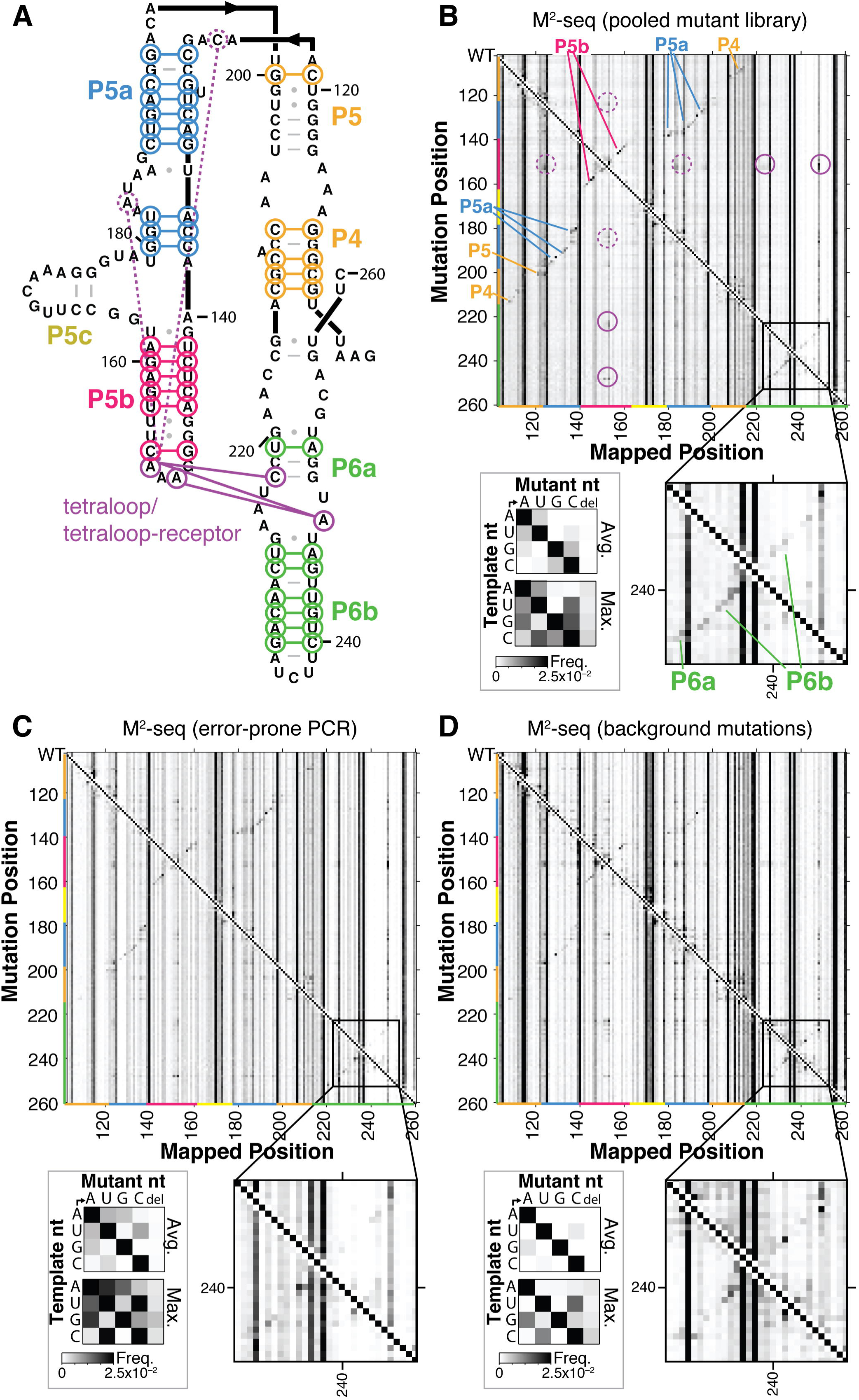
M2-seq on the P4-P6 domain of *Tetrahymena* group I ribozyme. (A) Secondary structure diagram of P4-P6. (B-D) Two-dimensional datasets from M2-seq on pooled mutate-and-map library (B), on RNA with mutations installed during error-prone PCR of DNA template (C), and with no intentionally installed mutations (D), all probed with dimethyl sulfate mapping. Each row gives frequencies of observing mutations at every position given a mutation at the row position, as indicated by the strong diagonal, top left to bottom right. In (B-D), black-lined outsets highlight M2-seq signals in the P6a-P6b region; gray-lined outsets show average and maximum observed frequencies of each type of mutation in control RNA samples without DMS treatment. In (A-B), colored lines and labels mark correspondence of structure and map signals for Watson-Crick helices, the tetraloop/tetraloop receptor contact (solid purple), and exposure of tetraloop from mutations outside its receptor (dashed purple).

Analysis of the mutational spectrum in the no-DMS samples confirmed that we had introduced the desired sequence changes at the level of ~1/158 (gray-lined outset, Fig. 2A). Furthermore, as expected, the M2-seq data (Fig. 2B) exhibit strong signals for structural elements, consistent with prior mutate-and-map data based on capillary electrophoresis (CE, Fig. S1). For example, M2-seq signals marking the pair C228/G246 and other pairs in the P6b helix create a visible cross-diagonal, as in prior CE data (black-lined outset, Fig. 2B). Base pairs for P4 and P5 (orange in Fig. 2B), P5a (blue), P5b (red), and P6a-b (green) are clearly visible and agree with crystallographic analysis of the RNA (Fig. 2A). Similarly, punctate signals reflecting the tetraloop/tetraloop receptor (TL/TLR) tertiary contact, such as between A153 and C223, also appear in both datasets. Short helices that were not observed in CE-based mutate-and-map measurements, such as the P5c helix (SI Fig. S1), also did not give extended cross-diagonal stripes in the M2-seq data. As expected, the no-DMS control samples did not show M2-seq signal, and consisted primarily of a uniform 1D background (Fig. S2A).

We further tested that separate preparation of mutants was not necessary to give clear M2-seq signals of base pairs. We used error-prone PCR to generate the DNA templates for RNA transcription, giving mutations at a mean frequency of ~0.5% and mostly involving U-to-C, C-to-U, A-to-G, and G-to-A transitions (gray-lined outset, Fig. 2C), as expected (19). Despite having a different mutational spectrum and giving signals at different specific base pairs, we observed M2-seq signals for the same helical elements as in the pooled single mutant library experiment as well as for the TL/TLR tertiary contact (Fig. 2C; fine differences are better visible in black-lined magnification outsets of P6a-P6b region). The use of error-prone PCR simplified the protocol: every step of the M2-seq experiment, from DNA synthesis to final reverse transcription and sequencing, could be carried out in a single tube.

We also observed M2-seq signal in samples without mutations intentionally installed during error-prone PCR (Fig. 2D). We had previously noted this pattern in published sequencing data for high-DMS-modified P4-P6 RNA (SI Fig. S3) (2) and speculated that DMS methylation of the N1 and N3 atoms of G and U residues, respectively, could disrupt Watson-Crick base-pairing, expose C and A partners, respectively, and produce a two-dimensional signal (2, 20). Paradoxically, however, the modification reaction pH of 7.0 is too low to cause significant deprotonation at these atoms to allow DMS methylation to occur (<10^−4^ modification rate expected under our conditions). Furthermore, when we applied the no-mutation method to another large, highly structured RNA, the *D. iridis* GIR1 lariat-capping ribozyme, we observed no clear cross-diagonal stripes corresponding to long-range RNA base pairs (Figure 3A).

**Figure 3.**
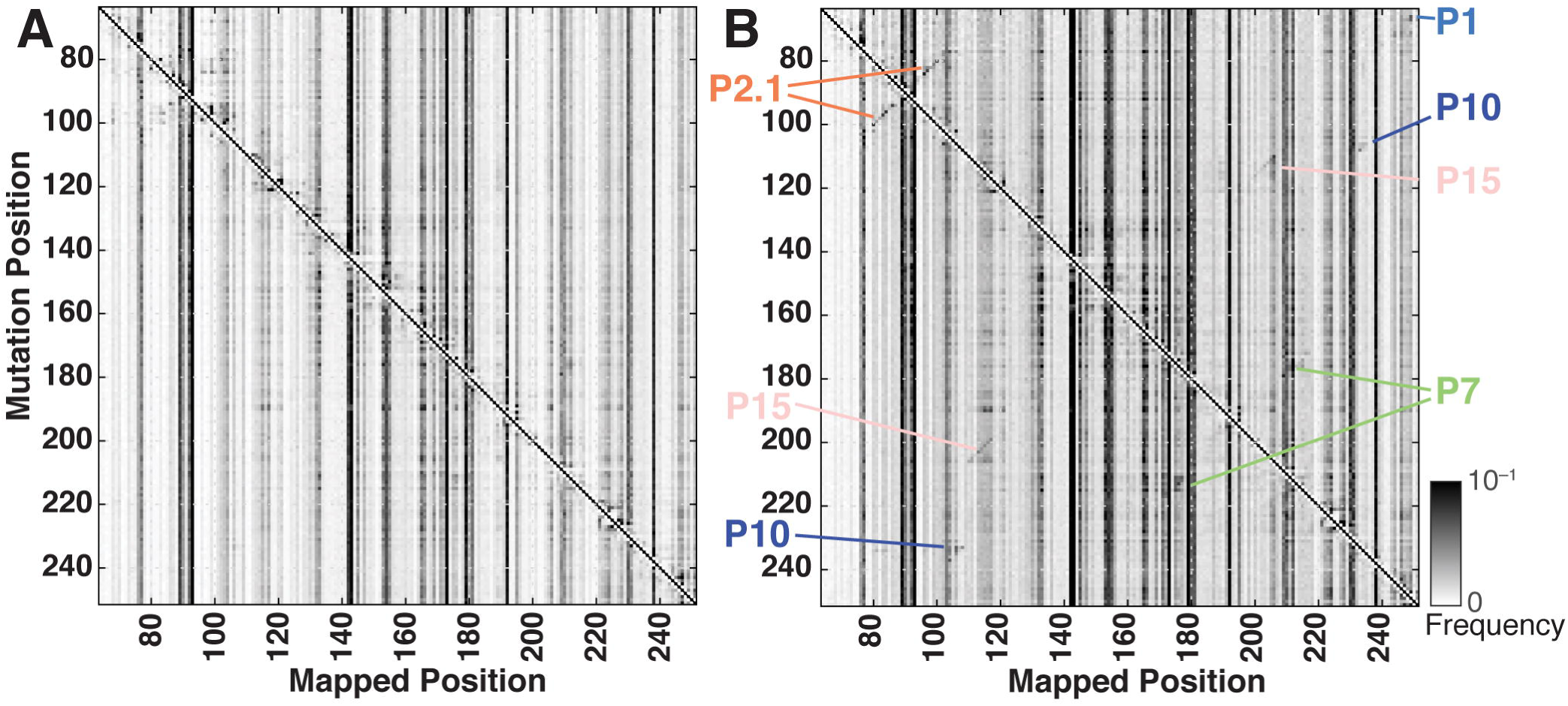
M2-seq on the GIR1 lariat-capping ribozyme requires seeded mutations. M2-seq maps for GIR1 ribozyme prepared (A) without any intentionally installed mutations and (B) from templates seeded with mutations through error-prone PCR. Colored text labels indicate helices for which cross-diagonal helix signatures become visible and detectable by M2-net.

### Mechanism of ‘background’ RNA base pair signals

To understand if M2-seq signals could be enhanced for the GIR1 ribozyme and other RNAs, we carried out extensive experiments to understand the mechanism for the signal in the P4-P6 RNA, varying transcription templates, purification, and modification conditions. A complete description of this work is given in SI Results and SI Figures S4-S7; a short summary follows. Briefly, we were able to discriminate between two models for how M2-seq signals might arise without intentionally pre-installed mutations. In a ‘double DMS hit’ model noted above, these Watson-Crick base pair signals are due to rare DMS modifications (~10 ^−3^ per nucleotide) that occur at transiently deprotonated U/G nucleotides, resulting in – or caused by – DMS modification at partner A/C nucleotides (2, 20). In an alternative ‘accidental mutation– model, the signals are due to background mutations (also up to ~10^−3^ per nucleotide) introduced as errors during DNA and RNA synthesis. In the folded RNA, these mutations would expose their structural partners to DMS, as with standard mutate-and-map methods.

Favoring the accidental mutation model, differences in the M2-seq signal with different DNA preparations (PCR assembly of oligonucleotides, PCR from a plasmid stock, and synthesis in other labs; Fig. 2 and SI Fig. S3) implicated background mutations introduced in the different DNA synthesis methods, which were then confirmed by sequencing the DNA templates used for those M2-seq experiments (Figs. S4 and S5). Additional M2-seq base pair signals were traced to transitions introduced during RNA synthesis by T7 RNA polymerase and confirmed by direct sequencing of the RNA before DMS modification (gray-framed outsets, Figs. 2, S4, and S5). Disfavoring the double DMS model, increasing the pH, which should enhance transient deprotonation of U/G and subsequent DMS modification, did not increase the M2-seq base pairing signal except at high pH. At pH 10.0, a different, less precise signal was observed (Fig. S6). Finally, DMS dose-response measurements revealed linear dependence of the Watson-Crick base pair signals with DMS dose, as predicted by the ‘accidental mutation’ model but not the ‘double DMS hit’ model, which predicts a quadratic dependence on DMS dose (Fig. S7).

Taken together, these studies traced the mechanism of direct base pair detection in DMS experiments to the occurrence of accidental mutations during DNA and RNA synthesis and not to double DMS hits. Because these mutations occur in a heterogeneous and non-controlled manner throughout the RNA molecule, they only allow detection of Watson-Crick pairs in special molecules with particular preparations. We therefore favored using error-prone PCR to seed in mutations more uniformly across transcripts. For example, in the case of the GIR1 lariat-capping ribozyme, M2-seq signals highlighting most of the RNA’s helices became visible when templates were prepared with error-prone PCR (Figure 3B). Even for the P4-P6 RNA, use of error-prone PCR allowed M2-seq detection of the P4-P6 helices with nearly an order of magnitude fewer sequencing reads than a protocol relying only on accidental mutations (SI Fig. S8).

### Automated detection of helices across diverse RNA structures

After testing the mechanism of the M2-seq signal, we evaluated the general applicability of the method across diverse structured RNA molecules. We chose several RNAs that have challenged prior structure modeling efforts: the P4-P6 RNA, the catalytic domain of RNase P, and the thiamine pyrophosphate riboswitch aptamer, which were the three test cases for an earlier RING-MaP study (20, 21); and the GIR1 ribozyme, riboswitch aptamers for adenosylcobalamin (AdoCBL) and cyclic-diAMP, and an Xrn1-exonuclease-resistant domain from the Zika virus, four targets of the RNA-puzzle community wide trials whose secondary structures were particularly challenging for most groups to model (12, 13, 23). M2-seq gave visually apparent signals for helices in all of these cases (Fig. 4 and SI Fig. S9). These helices included long-range interactions connecting the most sequence-distal ends of RNA (P1 in the GIR1 and RNAse P molecules), pseudoknots (P7 in GIR1; P2 in RNase P), long helices involved in tertiary contacts (9-bp P5b in P4-P6; 10-bp P8 in the AdoCBL riboswitch) and short helices (P3 in the TPP riboswitch). These signals were particularly apparent when we displayed maps of Z-scores, which measure how much the DMS signal at each nucleotide is enhanced over the mean at that position across all mutant variants, normalized to the standard deviation at that position. The quality of these data led us to revisit automated Z-score-based helix detection methods developed in early work on the mutate-and-map method (24, 25). Indeed, we discovered that our visual analysis could be automatically reproduced by a simple pipeline of Z-score estimation, a convolutional filter highlighting ‘cross-diagonal’ stripes, data symmetrization, and a filter for each nucleotide having at most one partner (SI Methods; colored annotations in Fig. 4). We called this analysis M2-net, due to its similarity to multi-layer convolutional neural nets that are now in wide use for image classification (26).

**Figure 4.**
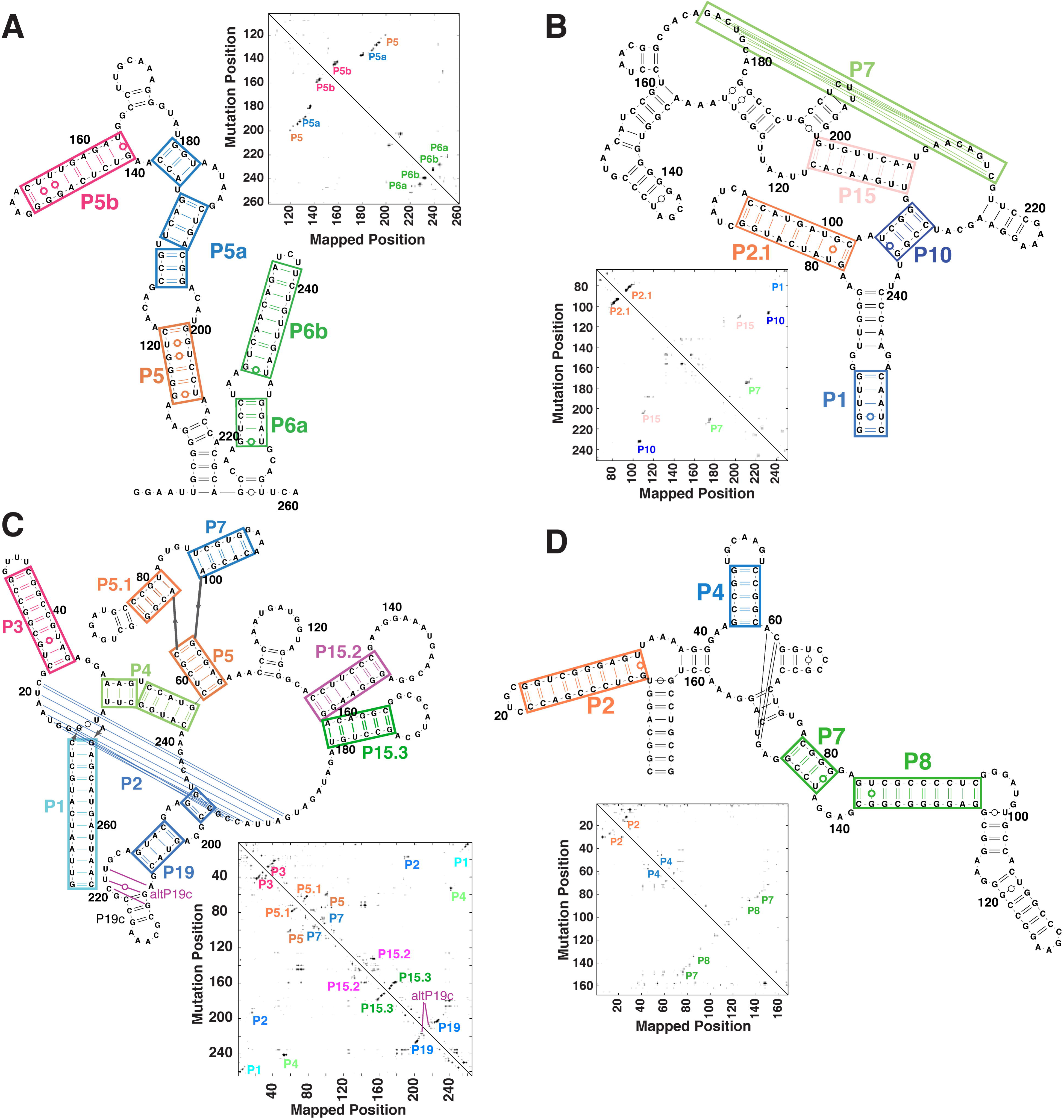
M2-seq recovers helices across diverse RNA folds. Each panel shows crystallographic secondary structures and Z-score-transformed maps (square graphs) with colored labels (on both display items) marking helices and multi-helix domains automatically identified by M2-net analysis. Differences in edge base pairs are not shown. Data sets are: (A) P4-P6 domain of *Tetrahymena* ribozyme (background mutations), (B) GIR1 lariat-capping ribozyme (RNA-puzzle 5; error-prone PCR), (C) Ribonuclease P catalytic domain (background mutations), (D) adenosylcobalamin riboswitch aptamer (RNA-puzzle 5; error-prone PCR). Data for three additional RNAs of smaller length are given in SI Fig. S9. SI Table S3 compiles modeled structures.

M2-net detected 34 of 60 helices with length greater than 2 in these RNAs (Table 1). 33 of these 34 helices matched the crystallographic or conventional structure available in the literature, and none of these cases involved register shifts that have been problems in prior methods (2, 7). Despite the observation of other weak signals in these data that do not correspond to helices (Fig. 4), M2-net detects only a single false-positive, a new alt-P19 helix predicted for the catalytic domain of RNase P that disagrees with the tip of the P19 domain presumed in the conventional secondary structure of this molecule (27). The region including these helices has not been directly visualized by crystallographic analysis (27), and we speculate that this RNA domain may interconvert between P19 and alt-P19 in solution.

**TABLE 1.** Recovery of helices across seven complex RNA folds from M2-seq data.

| RNA | Number of Helices<sup>a</sup> | ShapeKnots + DMS + M2<sup>b</sup> |  | M2-net |  |
| --- | --- | --- | --- | --- | --- |
|  |  | TP | FP | TP | FP |
| **P4-P6 domain** | 8 | 8 | 1 | 7 | 0 |
| **GIR1 ribozyme** | 11 | 11 | 0 | 5 | 0 |
| **RNase P C domain** | 13 | 11 | 1 | 10 | 1 |
| **AdoCbl riboswitch** | 10 | 10 | 0 | 4 | 0 |
| **ydaO riboswitch** | 7 | 6 | 1 | 2 | 0 |
| **Zika xrRNA** | 5 | 4 | 1 | 2 | 0 |
| **TPP riboswitch** | 6 | 6 | 0 | 3 | 0 |
| **Total** | 60 | 56 | 4 | 33 | 1 |
| **False negative rate (%)** |  | 6.7 |  | 45.0 |  |
| **False positive rate (%)** |  | 6.7 |  | 2.9 |  |
| **Sensitivity (%)** |  | 93.3 |  | 55.0 |  |
| **PPV (%)** |  | 93.3 |  | 97.1 |  |

<sup>a</sup> Helices with length greater than 2 Watson-Crick (or G•U wobble) base pairs.
<sup>b</sup> Use of one-dimensional DMS data to guide folding through energy bonuses (Cordero et al., Biochemistry. 2012;51(36):7037-9) and Z-scores derived from two-dimensional M2-seq experiments, applied as in (Kladwang et al., Nature Chemistry. 2011;3:954-62).

In prior work, we and others have used the RNAstructure free energy minimization software, guided by mutate-and-map or conventional one-dimensional chemical mapping data, to ‘fill in’ helices not directly detected by experiments (12, 20, 25). In our M2-seq benchmark, the ShapeKnots algorithm of RNAstructure guided by the M2-seq and one-dimensional DMS data indeed increases the number of recovered crystallographic helices from 34 to 56 (out of 60 helices; 93% sensitivity). However, the higher recovery is at the expense of more false positives: 4 out of a total of 60 predicted helices are incorrect (Table 1; SI Table S1 also includes modeling without pseudoknots and without DMS data). The resulting false discovery rate (7%) is similar to the rate seen in prior mutate-and-map benchmarks (16). For new RNAs where false positives would require expensive subsequent experiments to falsify, M2-net (with a false positive rate < 5%) may be preferred over RNAstructure analysis.

### RNA base pair detection in *Xenopus* egg extract

The simplicity of the ‘one-pot’ M2-seq protocol and the positive predictive value of the M2-net analysis motivated us to test the method in a more complex biological environment than the *in vitro* folding conditions typically used in benchmarking new chemical mapping methods. We mixed the P4-P6 RNA into undiluted extract from metaphase-arrested *Xenopus* eggs, a widely used medium for reconstituting eukaryotic biological processes (28). The impact of this complex medium, compared to *in vitro* conditions, was apparent in a new modification signature that arose *in extracto* but not *in vitro*, even in the absence of DMS treatment: A’s across the transcript were mutated to G. These modifications likely reflect the activity of the ADAR enzyme, which targets adenosine near double-stranded RNA helices for deamination to inosine, which is, in turn, read out as guanosine by reverse transcriptase (29, 30). Even with the complexities of *Xenopus* egg extract environment, the two-dimensional M2-seq data gave unambiguous signals for the P4-P6 RNA secondary structure. While these signals were less visually clear than in our *in vitro* experiments (Fig. 5B), they became more apparent when the data were viewed as Z-score maps (Fig. 5C). Despite an increase in background, partially due to A-to-I mutations, M2-net detected 5 of the 8 helices of P4-P6 and no false positives (colored labels in Fig. 5C).

**Figure 5.**
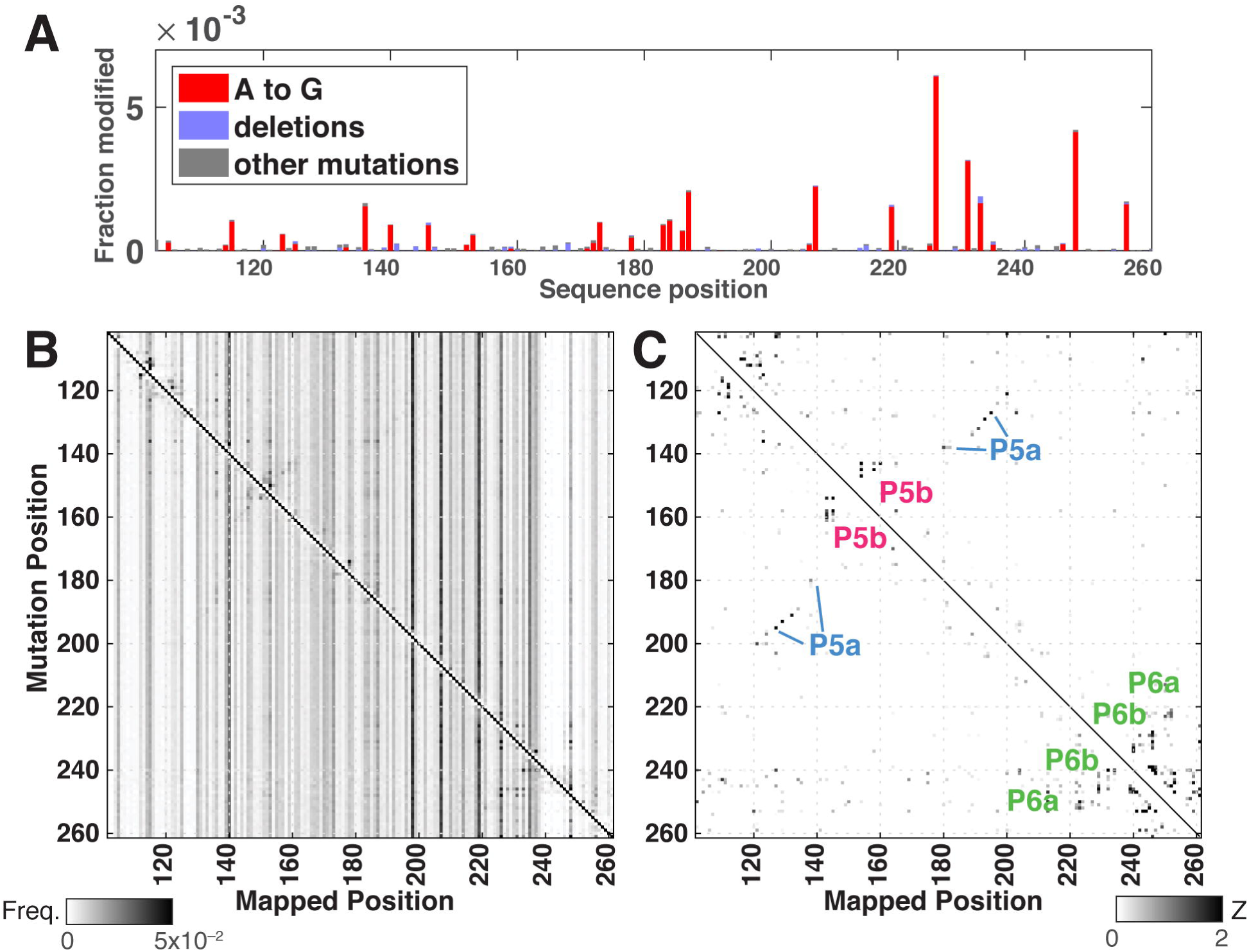
M2-seq detects P4-P6 RNA base pairs in *Xenopus* egg extract. Mutations across the P4-P6 transcript consistent with adenosine-to-inosine edits after exposure to undiluted *Xenopus* egg extract for 30 minutes (no DMS treatment); difference data with RNA incubated *in vitro* are shown. M2-seq data for P4-P6 RNA from templates prepared by error-prone PCR shown as (B) M2-seq map and then (C) transformed into Z-scores. Helix signatures automatically detected by M2-net are marked with colored labels.

## DISCUSSION

Rapid detection of base pairing partners in new non-coding RNAs has been difficult, requiring structural and biochemical techniques with low throughput, limited applicability, and/or poor predictive value. To address this challenge, we have introduced and tested a method called M2-seq (<u>m</u>utate-and-<u>m</u>ap read out through next-generation <u>seq</u>uencing). Mutations introduced at a low level (≲10^−3^) during DNA or RNA synthesis disrupt local structure in the folded RNA and expose interacting nucleotides to reaction with DMS. This mutation and a partner that becomes exposed to DMS methylation leave correlated imprints on single molecules, enabling readout through reverse trancription and next-generation sequencing. M2-seq permits precise detection of the major structural elements of classic model systems such as the 158-nucleotide P4-P6 domain of the *Tetrahymena* group I ribozyme and the 265-nucleotide *B. stearothermophilus* RNase P catalytic domain. M2-seq also reveals helices that have been difficult to detect or entirely missed in recent RNA-puzzles modeling for the GIR1 lariat-capping ribozyme, the adenosylcobalamin riboswitch, the *ydaO* cyclic diAMP riboswitch, and the Zika virus Xrn1-resistant genomic domain. Overall, the M2-seq data recover half of the helices in the tested RNAs with a low false positive rate (< 5%). Finally, the method enables pair-wise structure inference for the majority of helices of the P4-P6 RNA in *Xenopus* egg extract. To our knowledge, this is the first report of a biochemical technique enabling direct two-dimensional visualization of RNA base pair partners – as opposed to one-dimensional protections of uncertain origin – in a complex biological environment.

To supplement and automate simple visual inspection of M2-seq data, we have introduced the M2-net algorithm to infer helices from cross-diagonal signatures within the data, without bias from secondary structure modeling methods that attempt to minimize a computed free energy. The M2-net algorithm is expected to be particularly important for scenarios that are not appropriately modeled with energy minimization methods, such as cases involving non-trivial tertiary structure or multiple secondary structures, molecules with long lengths, or systems reconstituted in complex environments with protein binding partners or molecular machines that prevent the RNA from reaching equilibrium. Prior studies involving visual inspection of mutate-and-map data have correctly predicted tertiary contacts as well (12), and it will be important to test if M2-net can be expanded to inferring such 3D information.

The presented M2-seq protocol is immediately applicable to 250-nucleotide windows of lightly mutated RNAs introduced into complex biological environments. Synthetic long read sequencing or third-generation sequencing technologies may allow future studies to detect base pairings involving sequence separations longer than 250 nucleotides (31-33). In terms of seeding mutations, applications to viruses and other systems that involve high-error-rate RNA polymerases may obviate this step, but generally M2-seq in extracts, cells, and tissues will require transfecting DNA or RNA libraries that are prepared through error-prone PCR or other emerging techniques (32, 33). A faster and less biologically perturbing protocol would be enabled by a cell-permeable mutagen that could directly attack nucleotides initially sequestered inside RNA helices. While none of the routinely used chemical probes (e.g., DMS, SHAPE) appears appropriate, a large arsenal of mutagens remains to be tested for RNA structure mapping *in vivo* (34).

## METHODS

DMS mapping experiments on RNA were performed by modifying the RNA with DMS (170 mM final) in 10 mM MgCl_2_ and 300 mM Na-cacodylate (pH 7.0) for 6 min at 37 °C, followed by quenching with β-mercaptoethanol and purification with ethanol precipitation. Experiments with *Xenopus* egg extract replaced ethanol precipitation with purification by Trizol extraction and RNA Clean-and-Concentrator-5 columns (Zymo research). Reverse transcription was performed in conditions that lead to mutational readthrough at methylated nucleotides (SuperScript II and Mn^2+^), and sequencing libraries were prepared by PCR and sequenced on Illumina MiSeq instruments. ShapeMapper (35) was used to align sequencing reads to reference sequences and record mutations, and the results were converted to M2-seq data and mutation spectra using scripts available at https://github.com/DasLab/M2seq. Detailed descriptions of RNA preparation, modification experiments, map visualization, secondary structure modeling by M2-net and RNAstructure executables, and DMS dose-dependent mutation rate analysis are provided in the SI Methods.

## NOTE ADDED IN PROOF

Compensatory mutagenesis experiments carried out after paper review confirm the altP19c helix for RNase P shown in Fig. 4C; see SI Figure S10.

## ACKNOWLEDGMENTS

We thank P. Cordero and S. Tian for initial RNAstructure analyses of previously collected MaP data; the Herschlag lab for the pT7L-21 plasmid; B. French and A. Straight for the generous gift of *Xenopus* egg extract; and D. Mathews and R. Watson for incorporating extensions into RNAstructure software. We acknowledge funding from the National Institutes of Health (5 T32 GM007276 to C.Y.C.; R01 GM102519 to R.D.) and the Burroughs Wellcome Fund (CASI 1007326.01 to R.D.).

